# Robean: A Standalone Freeware for Automated Rodent Neurobehavioural Analysis with Integrated Tracking, Visualization, and Reporting

**DOI:** 10.64898/2026.07.11.737999

**Authors:** Vikas Mishra, Rajkumar Verma, Paruvathanahalli Siddalingam Rajinikanth, Ravinder K. Kaundal

## Abstract

Quantitative analysis of rodent behaviour is fundamental to neuroscience, preclinical drug discovery and neurotoxicology research. Although several commercial and open-source software packages are available for behavioural assessment, some are expensive, some require programming expertise, and some provide limited flexibility for user-defined experimental configurations. To address these limitations, we developed **Robean**, a freely available standalone software platform for automated rodent neurobehavioural analysis from both live camera feeds and pre-recorded videos.

Robean provides an intuitive graphical user interface that enables users to design experimental arenas, define custom analysis zones, perform spatial calibration, and automatically track rodent movement without requiring programming knowledge. The software currently supports automated analysis of three widely used behavioural paradigms: the Morris Water Maze, Elevated Plus Maze, and Open Field Test. Robean extracts behavioural metrics including escape latency, path efficiency, platform crossings, target quadrant preference, thigmotaxis, locomotor activity, zone occupancy, arm entries, and centre exploration specific to behavioural tests. In addition, the software generates trajectory maps, occupancy heatmaps, comma-separated value (CSV) datasets, comprehensive PDF reports, and batch study summaries for multiple experimental sessions.

Developed using open-source software technologies and distributed as a standalone freeware application, Robean provides an accessible and reproducible solution for behavioural neuroscience laboratories. Its modular architecture facilitates future integration of additional behavioural paradigms and analytical modules, making it a flexible platform for automated rodent behavioural assessment.

## 1. Introduction

Behavioural assessment of laboratory rodents is an indispensable component of preclinical drug discovery, neuroscience, behavioural, neurotoxicology, and neurological disease research. Experimental paradigms such as the Morris Water Maze (MWM), Elevated Plus Maze (EPM), and Open Field Test (OFT) are routinely employed to evaluate learning and memory, anxiety-like behaviour, locomotor activity, exploratory behaviour, and the efficacy of pharmacological or genetic interventions (Vorhees & Williams, 2014; Kodali et al, 2015; Acikgoz et al, 2022; Vorhees & Williams, 2024). Reliable quantification of behavioural parameters is therefore essential for generating reproducible and scientifically meaningful results.

Traditionally, behavioural scoring was performed manually by trained observers. Although manual scoring remains useful for certain specialised applications, it is inherently time-consuming, labour-intensive, and susceptible to observer bias and inter-rater variability. Automated video-based behavioural analysis has substantially improved the objectivity, reproducibility, and throughput of rodent behavioural studies, making computer-assisted tracking an integral part of modern behavioural neuroscience laboratories (Isik & Unal, 2023).

Several commercial software packages, including AnyMaze, EthoVision XT, SMART, and VideoMot, provide robust behavioural tracking and analysis capabilities across a variety of experimental paradigms (Noldus et al, 2001; Lim et al, 2023). However, their widespread adoption is frequently limited by high licensing costs, restricted customisation, proprietary algorithms, and dependence on commercial ecosystems. Conversely, numerous freely available and open-source tracking tools have been developed in recent years, yet many require programming expertise, provide limited graphical interfaces, or are designed primarily for a single behavioural paradigm. Consequently, researchers often need to combine multiple software tools or perform additional manual processing to obtain all behavioural metrics required for a study (Panaderio et al, 2021; Samson et al, 2015; Bello-Arroyo et al, 2018; Freeman et al, 2010; Forero et al, 2023).

To address these limitations, we developed Robean (Rodent Behavioural Analysis), a standalone free software platform for automated rodent neurobehavioural analysis. Robean has been designed to provide a user-friendly graphical interface while maintaining the flexibility expected from a freeware. The application supports both pre-recorded videos and live camera acquisition, enabling investigators to perform behavioural analysis without requiring programming knowledge or commercial software licenses.

Robean integrates multiple stages of behavioural analysis within a single application, including experimental arena creation, user-defined zone generation, spatial calibration, automated rodent tracking, behavioural metric extraction, trajectory visualisation, occupancy heatmap generation, data export, automated PDF reporting, and batch study summaries. The current implementation supports three of the most widely employed rodent behavioural paradigms: the Morris Water Maze, Elevated Plus Maze, and Open Field Test. In addition to single-experiment analysis, Robean supports automated batch processing of multiple behavioural recordings, enabling rapid generation of standardized behavioural reports for large experimental studies. Furthermore, its modular software architecture facilitates future incorporation of additional behavioural paradigms and analytical modules without requiring substantial modifications to the existing framework.

The objective of the present work is to introduce Robean as an accessible, reproducible, and freely available software platform for automated rodent behavioural analysis. By combining ease of use with comprehensive behavioural analysis and automated reporting, Robean provides researchers with a practical alternative to proprietary behavioural analysis systems. Its modular architecture also facilitates the future integration of additional behavioural paradigms and analytical modules, ensuring continued expansion of the platform.

## 2. Software workflow

Robean has been designed to provide a straightforward workflow that minimises user intervention while ensuring reproducible behavioural analysis. The complete analysis pipeline consists of nine sequential stages, beginning with video acquisition and concluding with automated report generation (Fig. 1).

**Figure 1.**
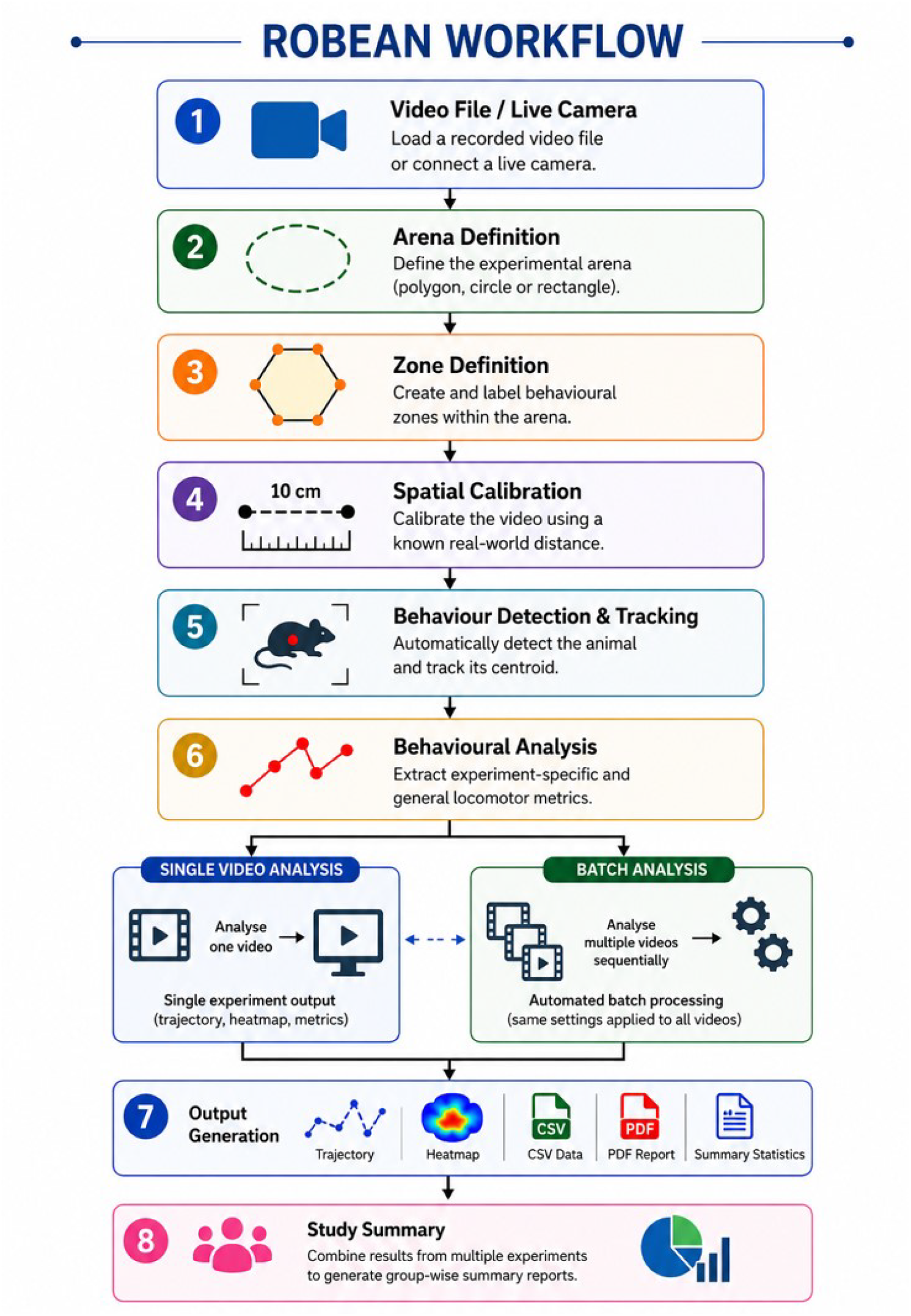
Overall workflow of Robean. The analysis pipeline begins with video input and ends with automatic generation of CSV files, PDF reports and study summaries.

### 2.1 Video acquisition

Behavioural analysis may be performed using either pre-recorded video files or live camera acquisition. Pre-recorded videos are optimised for MP4 format, while other commonly used formats are automatically converted before analysis. For live experiments, Robean supports multiple connected cameras, allowing users to select the desired imaging device.

### 2.2 Arena definition

The experimental arena is defined interactively using the graphical interface. Robean currently supports polygonal, circular, and rectangular arenas, enabling adaptation to different behavioural apparatuses. Previously created arenas can be saved and reloaded for future experiments, thereby improving experimental consistency across studies.

### 2.3 Zone definition

Behavioural analysis zones are created by drawing user-defined polygons within the arena. Each zone can be assigned a functional category according to the experimental paradigm, including platform, target quadrant, open arm, closed arm, centre, wall zone, periphery, or general regions of interest. Saved zone configurations can be reused across multiple experimental sessions.

### 2.4 Spatial calibration

To convert image measurements into real-world distances, users perform spatial calibration by selecting two reference points separated by a known physical distance. All locomotor parameters, including distance travelled, speed, and platform distance, are subsequently reported in calibrated physical units.

### 2.5 Behaviour detection and tracking

Following calibration, the animal is detected and tracked automatically throughout the recording. Robean continuously estimates the rodent centroid and updates behavioural information in real time. Movement trajectories, zone occupancy, locomotor statistics, and behavioural metrics are simultaneously computed during analysis.

### 2.6 Behavioural analysis

Experiment-specific behavioural parameters are calculated automatically according to the selected arena and zone configuration. General locomotor measurements, including distance travelled, average speed, movement trajectory, occupancy heatmap, and zone statistics, are also generated irrespective of the experimental paradigm.

### 2.7 Visualisation

During or after analysis, Robean allows users to visualise the complete movement trajectory and occupancy heatmap. These graphical representations facilitate rapid qualitative assessment of spatial exploration patterns and complement the quantitative behavioural metrics.

### 2.8 Data export

Following completion of analysis, behavioural data can be exported as comma-separated value (CSV) files for statistical analysis. Robean simultaneously generates PDF reports containing experiment details, behavioural metrics, trajectory images, occupancy heatmaps, and summary statistics.

### 2.9 Study summary and batch analysis

For experiments comprising multiple animals or treatment groups, Robean provides an automated Study Summary module that combines behavioural metrics from individual experiments into a single consolidated report. This feature simplifies comparison between experimental groups and reduces manual data compilation. Further for multiple studies, Robean also provides an automated Batch Analysis module. Using previously defined arena, zone, and calibration settings, multiple video files can be processed sequentially without further user intervention. Individual behavioural outputs are generated automatically for each recording, followed by an integrated study summary.

The overall workflow implemented in Robean enables behavioural experiments to be analysed using a consistent sequence of operations while minimising user-dependent variability. The integrated design eliminates the need to transfer data between multiple software packages and provides a unified environment for behavioural tracking, analysis, visualisation, and reporting.

## 3. Software requirements, architecture and implementation

Robean was developed as a standalone desktop application using the Python programming language and open-source scientific computing libraries. The graphical user interface (GUI) was implemented using the Qt framework, providing an intuitive, interactive environment for behavioural analysis without requiring programming expertise. Image acquisition and processing are performed using OpenCV, while data analysis, report generation, and file management rely exclusively on open-source Python libraries. The software is distributed as a self-contained executable, enabling installation and execution without requiring a separate Python environment.

### 3.1 System requirements

The current version of Robean is designed for 64-bit Microsoft Windows operating systems and has been tested on Windows 10 and Windows 11. Earlier versions of Windows (Windows 8.1 and below) are not supported. The software requires a minimum of an Intel Core i3 (8th Generation or later), AMD Ryzen 3 processor, or an equivalent CPU, although an Intel Core i5/i7 or AMD Ryzen 5/Ryzen 7 processor is recommended for improved performance during video processing. A minimum of 8 GB RAM is required, while 16 GB RAM is recommended for efficient analysis of large video datasets. Approximately 500 MB of free disk space is required for software installation, with additional storage depending on the size and number of video files and generated analysis reports.

### 3.2 Software architecture

The software follows a modular architecture in which individual components are responsible for independent stages of behavioural analysis. This design simplifies maintenance, facilitates future expansion, and allows new behavioural paradigms or analytical modules to be incorporated with minimal modifications to the existing framework (Fig. 2).

**Figure 2.**
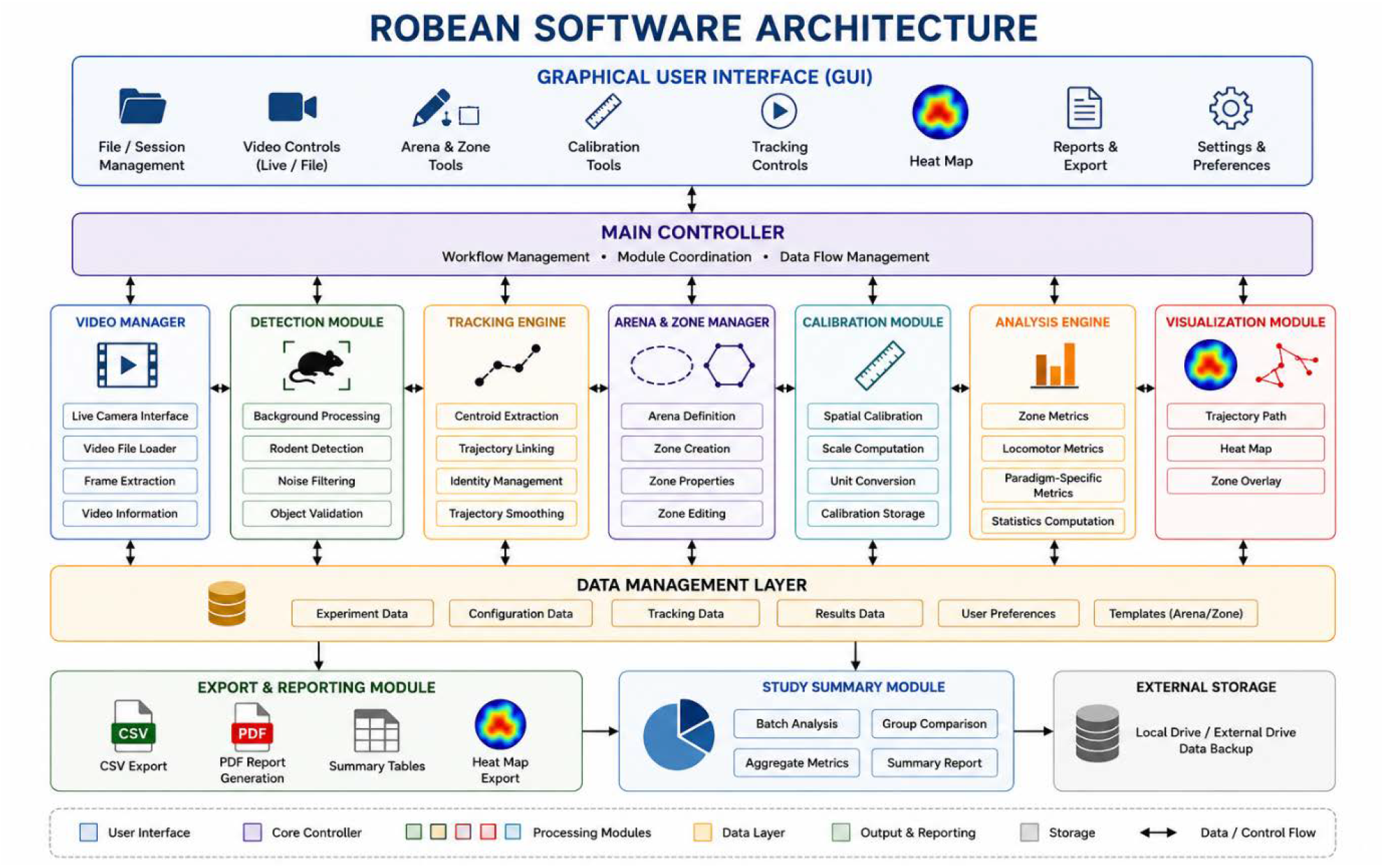
Software architecture of Robean. The system is organized into modular components that work together through the main controller and data management layer to deliver automated behavioural analysis and reporting.

The principal software modules include:

#### 3.2.1 Video acquisition module

The video acquisition module **s**upports both pre-recorded video files and real-time acquisition from USB or integrated cameras. Multiple cameras can be selected when available, allowing the software to be used for both offline and live behavioural experiments.

#### 3.2.2 Arena management module

This module provides interactive tools for constructing experimental arenas using polygonal, circular, or rectangular geometries. Previously created arenas can be saved, reloaded or reused across multiple experiments.

#### 3.2.3 Zone management module

The zone management module allows users to define an unlimited number of user-defined analysis zones within an experimental arena. The zones may be assigned various functional categories, including platform, target quadrant, open arm, closed arm, centre, wall zone, periphery, or general regions of interest, enabling automatic paradigm-specific behavioural analysis. Moreover, previously created zones can be saved, reloaded or reused across multiple experiments (Fig. 3).

**Figure 3.**
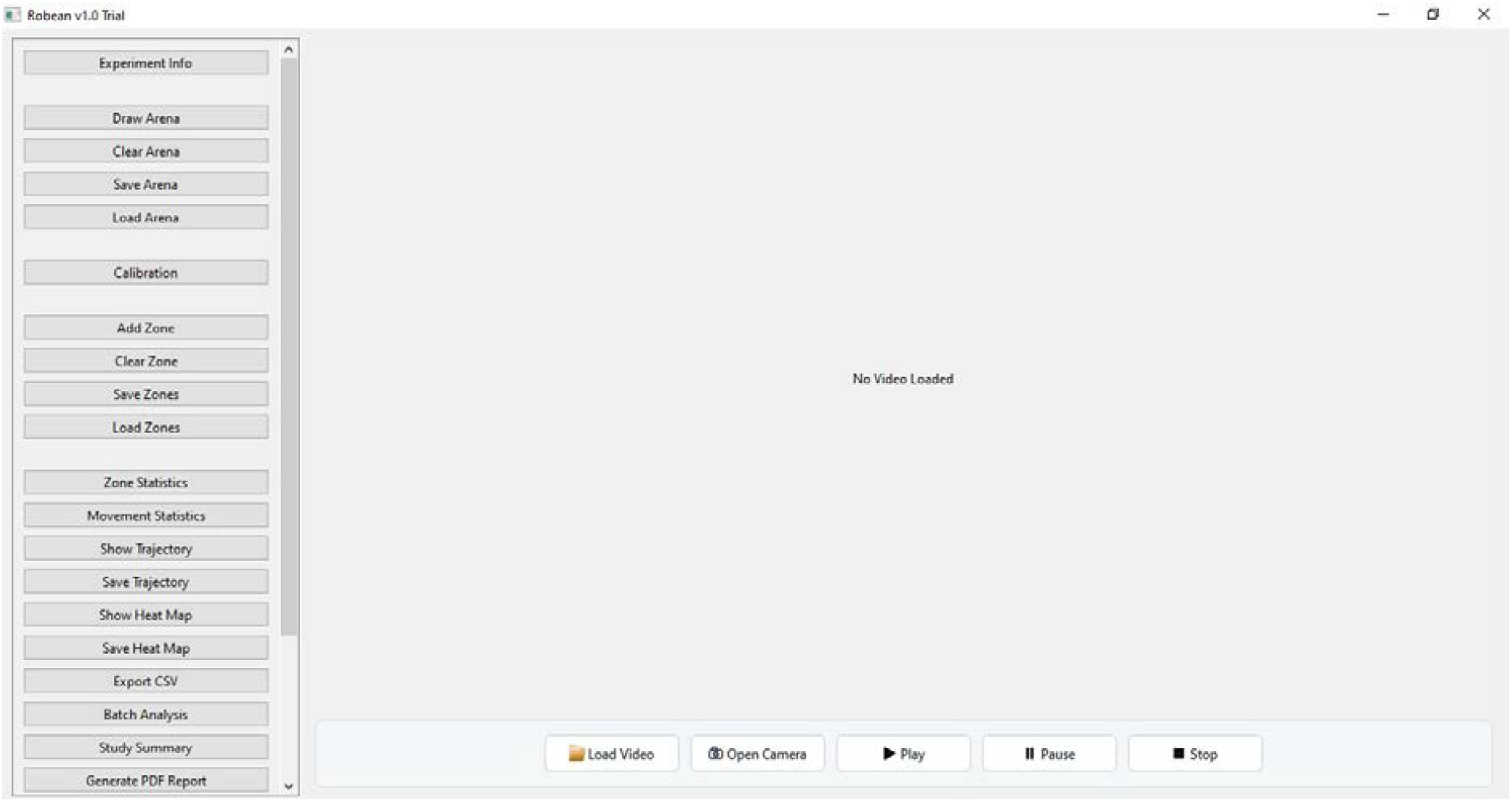
Main graphical user interface (GUI) of Robean. The interface integrates video loading and acquisition, arena and zone management, spatial calibration, behavioural analysis, and visualization of tracking results. It also supports trajectory and heat map generation, batch analysis, and export of results as CSV files or PDF reports.

#### 3.2.4 Calibration module

The calibration module converts image coordinates into real-world spatial measurements using user-defined reference distances, thereby allowing behavioural metrics such as distance travelled and average speed to be reported in physical units.

#### 3.2.5 Detection and tracking module

The detection and training module automatically detects and tracks the centroid of the experimental animal throughout the recording (Fig. 4). The software supports both dark- and light-coloured rodents and continuously updates movement trajectories, spatial occupancy, and behavioural statistics during analysis.

**Figure 4.**
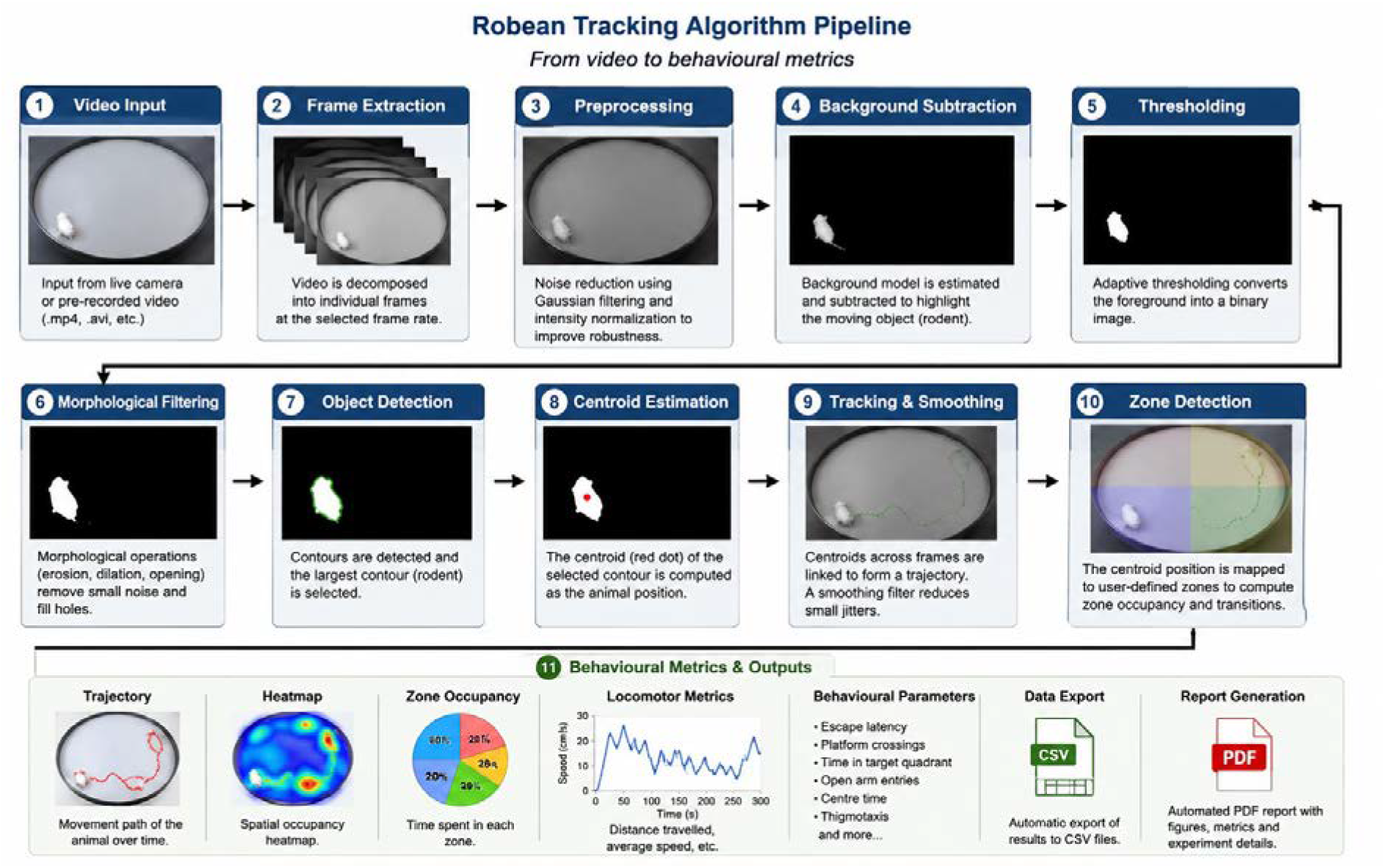
Robean Tracking Algorithm Pipeline: Robean uses a robust centroid based tracking approach combining image processing and spatial analysis to provide accurate, efficient and automated behavioural analysis for rodent experiments.

#### 3.2.6 Behavioural analysis module

The behavioural analysis module automatically computes behavioural parameters specific to the selected experimental paradigm. The current implementation supports automated analysis of the Morris Water Maze, Elevated Plus Maze, and Open Field Test while maintaining a common analysis framework applicable to future paradigms.

#### 3.2.7 Visualisation module

The visualisation module generates real-time movement trajectories and occupancy heatmaps, enabling rapid qualitative assessment of spatial exploration patterns in addition to quantitative behavioural metrics.

#### 3.2.8 Reporting module

Automatically exports behavioural results as comma-separated value (CSV) files and generates comprehensive PDF reports containing behavioural metrics, experiment information, graphical summaries, and spatial analyses. Multiple experiments can additionally be combined into study-level summary reports to facilitate statistical analysis across experimental groups. The reporting framework also incorporates an automated Batch Analysis workflow that sequentially analyses multiple behavioural recordings using identical analysis parameters, thereby improving throughput and ensuring analytical consistency across experimental datasets.

The modular organisation of Robean separates user interaction, video acquisition, behavioural tracking, statistical analysis, and report generation into independent software components. Such separation improves code maintainability, enhances software reliability, and provides a scalable framework for integrating additional behavioural paradigms, advanced tracking algorithms, and machine learning–based analytical methods in future releases.

## 4. Supported behavioural paradigms

The current version of Robean supports automated analysis of three widely employed rodent neurobehavioural paradigms: the Morris Water Maze (MWM), Elevated Plus Maze (EPM), and Open Field Test (OFT). Each paradigm has been implemented using a common tracking framework while providing experiment-specific behavioural metrics. User-defined arenas and analysis zones allow investigators to adapt the software to different experimental protocols without modifying the underlying tracking algorithm.

### 4.1 Morris water maze

The Morris Water Maze is one of the most extensively used paradigms for evaluating spatial learning and memory in rodents. During the acquisition phase, animals learn to locate a hidden escape platform using distal visual cues. Subsequently, probe trials assess memory retention by evaluating search behaviour after removal of the platform.

Robean automatically quantifies the principal behavioural endpoints assessed in Morris Water Maze studies, including escape latency, platform crossings, target quadrant occupancy, target quadrant preference, mean distance from the platform, path efficiency, thigmotaxis (wall-hugging behaviour), total distance travelled, and average swimming speed. These parameters are calculated automatically at the end of each experiment and can be exported as CSV files or PDF reports (Fig. 5).

**Figure 5.**
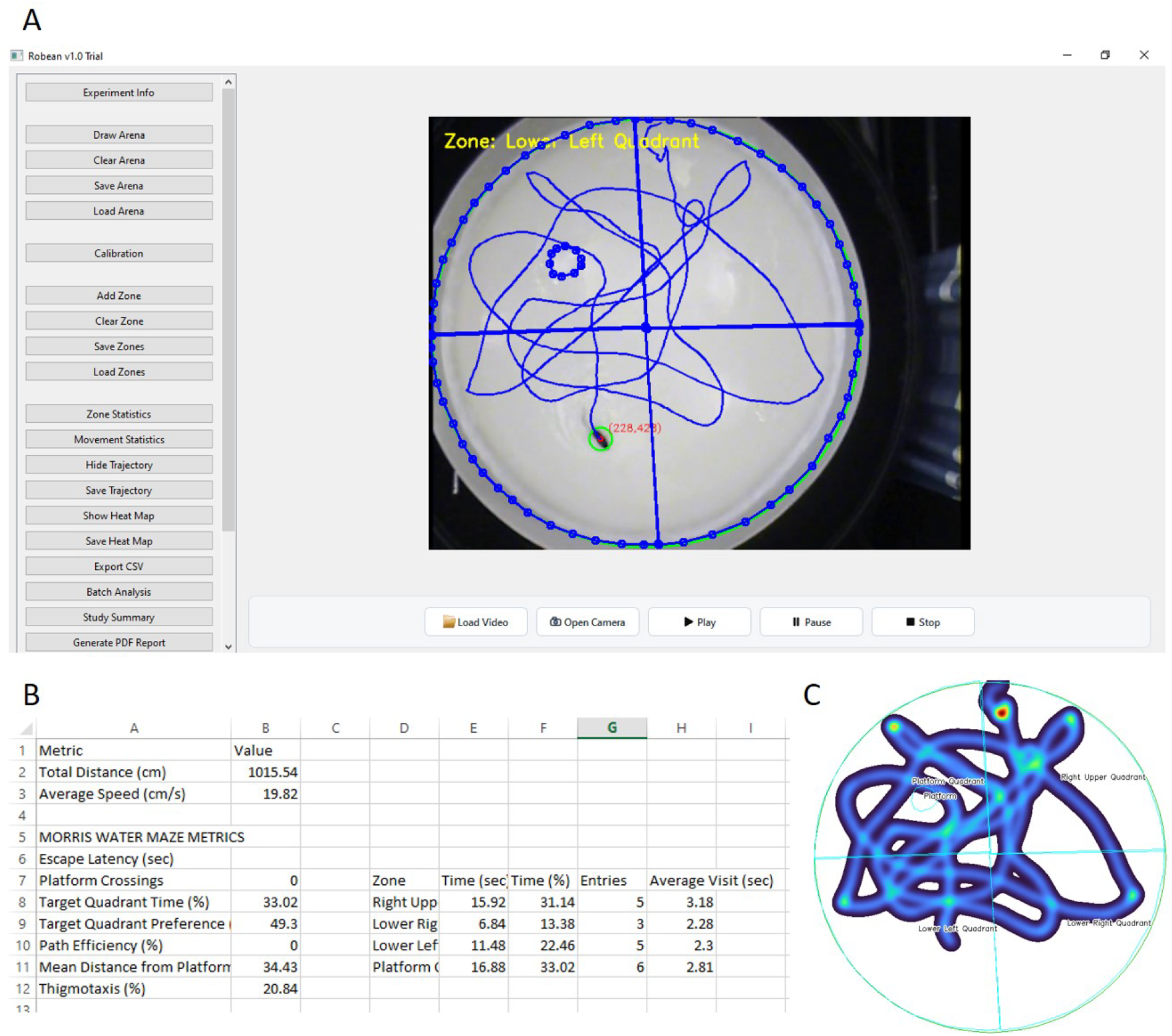
Representative analysis of a Morris Water Maze experiment using Robean. **(A)** Robean graphical user interface showing automatic detection of the rodent, centroid localization (red dot), user-defined arena and analysis zones, and reconstruction of the swimming trajectory in real time. **(B)** Representative output of behavioural parameters automatically generated by the software, including total distance travelled, average swimming speed, escape latency, platform crossings, target quadrant occupancy, target quadrant preference, path efficiency, mean distance from the platform, thigmotaxis, and zone-specific occupancy statistics. **(C)** Representative occupancy heat map illustrating the spatial distribution of the animal’s movement within the arena, enabling visualization of search strategy and quadrant preference.

### 4.2 Elevated plus maze

The Elevated Plus Maze is widely used for assessing anxiety-like behaviour in rodents. The apparatus consists of two open arms and two closed arms elevated above the floor. Rodents naturally avoid open spaces; therefore, increased exploration of the open arms is generally interpreted as reduced anxiety-like behaviour.

Robean automatically determines the number of open-arm entries and closed-arm entries, the percentage time spent in open arms, closed arms and central platform, open-arm preference, total distance travelled and average speed. These parameters are calculated automatically at the end of each experiment and can be saved as CSV files or PDF reports (Fig. 6).

**Figure 6.**
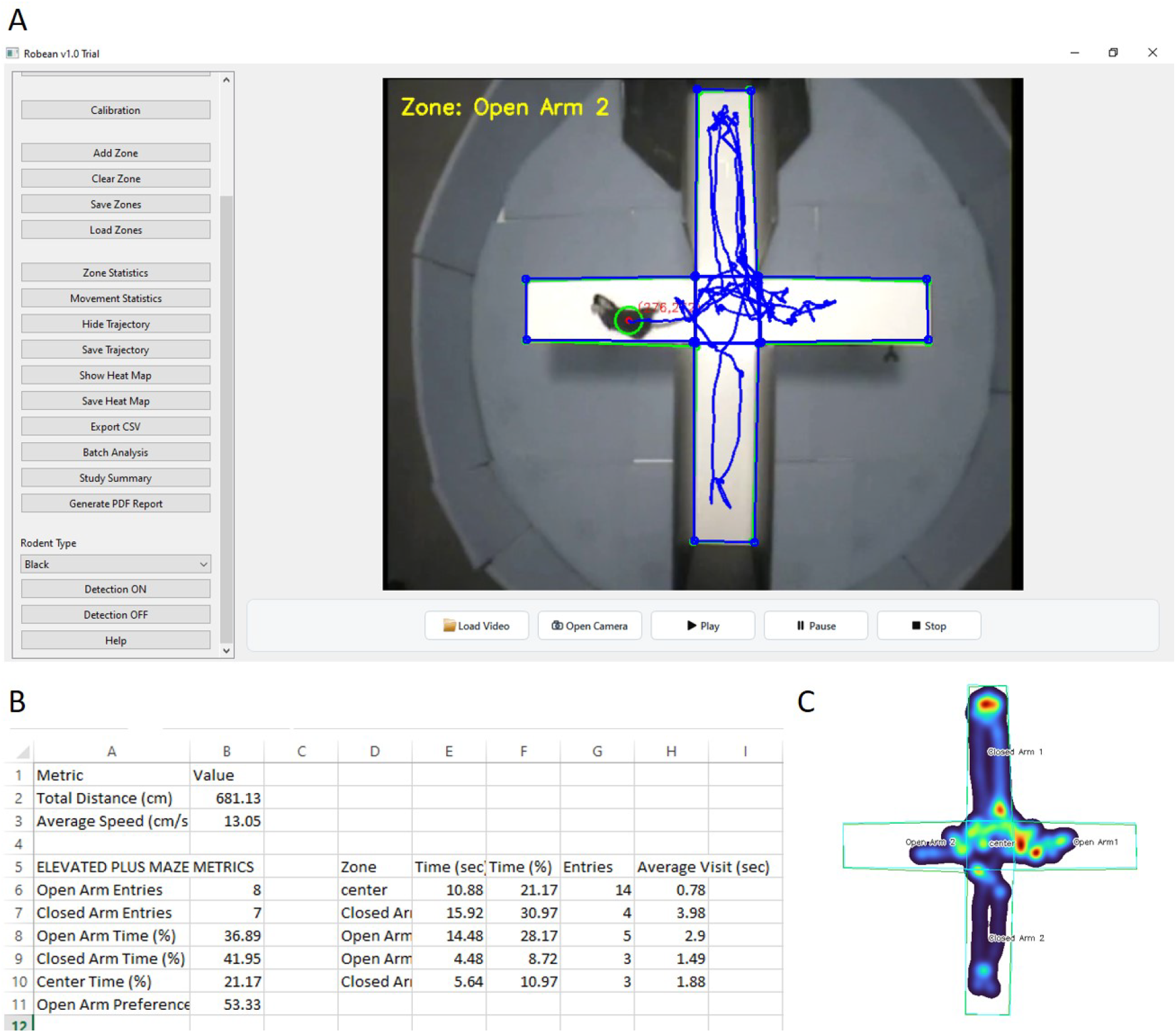
Representative Elevated Plus Maze analysis performed using Robean. **(A)** Robean graphical user interface showing automatic detection of the rodent, centroid localization (red dot), user-defined analysis zones corresponding to the open arms, closed arms, and central platform, and real-time reconstruction of the movement trajectory. **(B)** Representative spreadsheet output summarizing behavioural parameters automatically calculated by the software, including total distance travelled, average movement speed, open-arm entries, closed-arm entries, percentage time spent in the open arms, closed arms, and centre, open-arm preference, and zone-specific occupancy statistics. **(C)** Representative occupancy heat map illustrating the spatial distribution of the rodent’s movement within the maze, highlighting zone preference and exploratory behaviour.

Zone classification is entirely user-configurable, allowing the software to accommodate different maze dimensions and experimental designs.

### 4.3 Open field test

The Open Field Test is commonly employed to evaluate spontaneous locomotor activity, exploratory behaviour, and anxiety-related behaviour. Rodents generally prefer the peripheral region of the arena, whereas increased exploration of the central area is frequently interpreted as reduced anxiety-like behaviour.

Robean automatically quantifies centre occupancy, thigmotaxis (peripheral exploration), total distance travelled, and average locomotor speed. Because the arena and behavioural zones are user-defined, the software can readily accommodate open-field arenas of different sizes and geometries.

### 4.4 Common behavioural measurements

In addition to experiment-specific parameters, Robean continuously records general locomotor behaviour during all supported paradigms. These measurements include cumulative distance travelled and average speed. These common metrics provide additional behavioural information that may complement experiment-specific analyses.

### 5.0 Validation and performance evaluation

The current trial version of Robean was evaluated using pre-recorded behavioural videos obtained from standard rodent neurobehavioural experiments conducted during the first author’s postdoctoral research in the laboratory of Prof. Ashok K. Shetty at the Texas A&M Health Science Center, College of Medicine, Temple, Texas, USA. Software development and testing primarily focused on the Morris Water Maze (MWM) and Elevated Plus Maze (EPM), while the Open Field Test (OFT) module was subsequently incorporated using the same behavioural tracking framework.

Evaluation was performed under routine laboratory conditions using videos acquired from overhead cameras with fixed imaging geometry. Experimental arenas and behavioural zones were manually defined through the graphical interface prior to analysis, after which behavioural tracking and metric extraction were performed automatically. The software successfully tracked rodent movement throughout complete experimental sessions and generated behavioural outputs including movement trajectories, occupancy heatmaps, zone occupancy statistics, locomotor parameters, CSV datasets, and comprehensive PDF reports.

The Morris Water Maze module was evaluated for automatic estimation of escape latency, platform crossings, target quadrant occupancy, target quadrant preference, path efficiency, mean distance from the hidden platform, and thigmotaxis. Similarly, the Elevated Plus Maze module was assessed for automated quantification of open-arm entries, closed-arm entries, open-arm exploration time, closed-arm exploration time, centre occupancy, and open-arm preference.

The Batch Analysis module was evaluated using multiple prerecorded Morris Water Maze and Elevated Plus Maze videos. Consecutive recordings were processed automatically without manual intervention, producing behavioural metrics, trajectories, occupancy heatmaps, CSV datasets, PDF reports, and study summaries identical in format to those generated during individual analyses. These observations demonstrate the robustness and reproducibility of the batch-processing framework for large experimental datasets.

Performance testing demonstrated that Robean can analyse both prerecorded videos and live camera feeds through a unified graphical interface. In addition to behavioural metrics, the software automatically produces summary reports without requiring post-processing in external software. Arena configurations, zone definitions, and calibration settings can be saved and reused, thereby improving reproducibility across repeated experiments.

The present study was intended to demonstrate the functionality and practical applicability of Robean rather than to provide a comprehensive benchmarking study against commercial behavioural analysis systems. Formal validation against manual behavioural scoring and established commercial software platforms is planned for future releases. Such comparative evaluation will include assessment of tracking accuracy, agreement between automated and manual behavioural measurements, computational performance, and inter-software reproducibility across multiple neurobehavioural paradigms.

## 6. Discussion

Automated behavioural analysis has become an essential component of modern preclinical neuroscience and behavioural pharmacology. While several commercial systems provide highly accurate behavioural tracking, their widespread use is often limited by high acquisition costs, proprietary software ecosystems, and restricted opportunities for customization.

Conversely, many open-source behavioural analysis tools require programming expertise, installation of multiple software dependencies, or extensive post-processing before meaningful behavioural parameters can be obtained (Isik & Unal, 2023; Panadeiro et al, 2021). These practical limitations continue to present challenges for many academic laboratories, particularly those operating with limited financial or computational resources.

Robean was developed to address these challenges by providing a standalone freeware application that combines behavioural tracking, quantitative analysis, visualization, and automated report generation within a single graphical environment. The software has been designed to minimise user intervention while maintaining sufficient flexibility to accommodate different experimental protocols through user-defined arenas, behavioural zones, and calibration procedures. By integrating video acquisition, behavioural tracking, trajectory visualization, occupancy heatmap generation, CSV export, PDF reporting, and study-level summaries into a unified workflow, Robean substantially reduces the time required for behavioural data processing and eliminates the need for multiple independent software packages. An additional practical advantage is the integrated Batch Analysis module, which enables automatic processing of multiple behavioural recordings using identical experimental settings. This considerably reduces analysis time for large-scale studies while improving consistency by eliminating repetitive manual operations between individual recordings.

One of the principal strengths of Robean is its versatility. The current implementation supports three widely used behavioural paradigms—Morris Water Maze, Elevated Plus Maze, and Open Field Test—using a common analytical framework. This modular architecture enables the software to compute both paradigm-specific behavioural endpoints and general locomotor parameters simultaneously. The same tracking engine can therefore be extended to additional behavioural assays without fundamental modifications to the software architecture.

Another important feature of Robean is its support for both prerecorded video analysis and live camera acquisition. Real-time behavioural monitoring enables investigators to evaluate experiments as they are performed while preserving the ability to analyse archived recordings using identical analytical procedures. Furthermore, reusable arena and zone templates improve experimental reproducibility across repeated studies and reduce operator-dependent variability.

Future development of Robean will focus on expanding the range of supported behavioural paradigms, improving detection robustness using modern computer vision and deep-learning approaches, increasing compatibility with diverse camera systems, and incorporating additional analytical modules. Planned behavioural paradigms include Barnes Maze, Y-Maze, Novel Object Recognition, Light/Dark Box, Social Interaction Test, Radial Arm Maze, and other commonly employed neurobehavioural assays. Additional developments will include enhanced batch analysis, advanced statistical summaries, and support for user-defined behavioural plugins.

In conclusion, Robean provides a practical, flexible, and freely available solution for automated rodent neurobehavioural analysis. Its integrated workflow, intuitive graphical interface, comprehensive reporting capabilities, and support for multiple behavioural paradigms make it a valuable analytical tool for behavioural neuroscience, pharmacology, and biomedical research laboratories. Continued development and validation are expected to further enhance its applicability across a broad range of experimental paradigms.

## 7. Limitation

Although Robean demonstrated reliable performance during software development and laboratory evaluation, several limitations remain. The current tracking algorithm primarily relies on centroid-based image processing and therefore may be affected by poor illumination, excessive shadows, reflections, or substantial occlusion of the experimental animal. Camera performance may also vary depending on the imaging hardware and operating system. In addition, formal benchmarking against established commercial behavioural analysis software and manual behavioural scoring has not yet been completed and will be addressed in future validation studies.

## 8. Software availability

Robean is distributed as a standalone freeware application for Microsoft Windows (Windows 10 or later). The software can be installed without requiring a separate Python environment or additional third-party dependencies. The freeware version is intended for academic research and educational purposes. Future versions, software updates, user documentation, and example datasets will be made available through the official project webpage. The Robean software installer and accompanying documentation are freely available for academic and research use at: https://sourceforge.net/projects/robean/files/

## 9. Conclusion and future directions

Robean is a standalone freeware software platform developed for automated rodent neurobehavioural analysis from both prerecorded videos and live camera acquisition. By integrating behavioural tracking, arena and zone management, spatial calibration, automated metric extraction, trajectory visualization, occupancy heatmaps, CSV export, PDF reporting, and study-level summaries into a single graphical interface, Robean simplifies behavioural data analysis while improving experimental reproducibility.

The current implementation supports automated analysis of the Morris Water Maze, Elevated Plus Maze, and Open Field Test and provides a flexible architecture that can accommodate additional behavioural paradigms in future releases. Robean therefore represents an accessible and practical alternative for behavioural neuroscience laboratories seeking an integrated behavioural analysis platform without the financial barriers associated with commercial software. Future releases of Robean will focus on expanding behavioural paradigm support, improving tracking robustness using pose estimation and artificial intelligence techniques, and enhancing compatibility with a wider range of imaging hardware.

## Acknowledgements

The authors gratefully acknowledge Babasaheb Bhimrao Ambedkar University, Lucknow, India, for providing the academic environment and facilities necessary for the development and evaluation of Robean. The authors also acknowledge that the behavioural videos used in Figures 5 and 6 were generated during the postdoctoral research of Dr. Vikas Mishra in the laboratory of Prof. Ashok K. Shetty at the Texas A&M Health Science Center, College of Medicine, Temple, Texas, USA.

## Funding

This work received no specific grant from any funding agency in the public, commercial, or not-for-profit sectors.

## Conflict of Interest

The authors declare that they have no competing financial or non-financial interests related to this work.

## Data availability

The behavioural videos used during software development and evaluation are available from the corresponding author upon reasonable request subject to institutional policies.

The Robean software installer and accompanying documentation are freely available for academic and research use at: https://sourceforge.net/projects/robean/files/

## Author contributions

**Vikas Mishra:** Conceptualization, software design, software development, validation, data analysis, visualization, manuscript preparation, and project supervision.

**Raj Kumar Verma:** Software evaluation, behavioural validation, manuscript review, and scientific input.

**Paruvathanahalli Siddalingam Rajinikanth:** Software evaluation, manuscript review, and scientific input.

**Ravinder Kaundal:** Software evaluation, behavioural validation, manuscript review, and scientific input.

All authors reviewed and approved the final manuscript.

## AI acknowledgement

The authors used ChatGPT (OpenAI) as a software development and assistance tool during the design, implementation, debugging, and language editing of the Robean software. All scientific decisions, software validation, interpretation of results, and manuscript preparation were performed by the authors, who take full responsibility for the final work.

## Notes

https://sourceforge.net/projects/robean/files/

### Competing Interest Statement

The authors have declared no competing interest.

